# A spark of 3D revisualization: new method for re-exploring segmented data

**DOI:** 10.1101/2020.08.01.222869

**Authors:** Yuzhi Hu, A. Limaye, Jing Lu

## Abstract

3D scientific visualization is a popular non-destructive investigation tool, however current imaging processing and 3D visualization software has compatibility barriers which make replicability and reproducibility in research difficult. To solve this, we developed a new revisualization method and demonstrated four case studies using three mainstream image processing and 3D visualization software. Our method offers interchangeability amongst current image processing and 3D visualization software.

---

Nowadays, 3D scientific visualization, becomes a common solution for many different purposes^1–4^ and one of the most useful tools for research investigations, revealing hidden information behind the data^5–8^. For the last two decades, researchers in the 3D scientific visualization field have been experimenting with new ways to more accurately visualize medical and scientific data-sets^3,9–17^. In the current workflow of 3D scientific visualization process, 3D rendered results can be generated using combinations of image processing, segmentation, and rendering techniques^18–20^. Current image processing and 3D visualization software lack interchangeability and contain natural barriers due to compatibility (e.g. format protection between different software), accessibility (e.g. different researchers use different software so manageability of data sharing between collaborated parties is limited) and affordability (e.g. students may not afford the license for commercial software). The vast amount of time taken to master each software and the complexity behind different algorithms or approaches, are also a concern. These barriers weaken the achievability of the transparency, replicability and reproducibility of scientific research; damage the validation of communications between collaboration of institutions and restrict the authenticity assessment of a research finding, a proposed solution or a hypothesis.

It is a trend that more publishers and journals start requesting authors to upload sufficient data to support their discoveries and ensure published findings are authentic^14,21–24^. A common solution is uploading surface mesh data (i.e. polygon mesh, for example Stereolithography Format (. stl) or Polygon File Format (.ply)) of the segmented results of the research object. However, this approach still restricts the capability of re-investigation by peers as surface mesh data only contains external information. A more efficient and resource-friendly way is to use the segmented volume data, which contains both internal and external information together with the original volumetric data, however, as mentioned above, due to current restrictions and barriers, this is not currently feasible.

Most mainstream image processing and 3D visualization software (e.g. Mimics, Avizo, VG Studio) can input and output Digital Imaging and Communications in Medicine (DICOM). DICOM is well known in medicine for being exchangeable between any two entities in biomedical imaging and analysis software^25^. In medicine, segmented DICOM data from a scan (e.g. ultrasound or MRI) can be used to re-analysis segmented regions of interest using other software for diagnostics^26,27^. Thus, we developed volume exploration and presentation software *Drishti* to solve the current barriers in image processing and 3D visualization software, by implementing an ability to include both single and multiple DICOM directories of segmented volumetric data from other 3D visualization software. To the best of our knowledge, this method conquers current compatibility and interchangeability constraints among 3D visualization software.

By testing this new revisualization method, we use four segmented volumetric data from three mainstream 3D visualization software - Mimics, VG Studio and Avizo. The four segmented volume data have been saved as single (VG Studio) and multiple DICOM (Mimics and Avizo) directories, and then imported into *Drishti* to transform and converted into processed volume formats which *Drishti* can read and process. After proceeded to the processed volume format (i.e. .pvl.nc), segmented volume data can be explored and rendered in *Drishti* (Fig.1 and SI Figs.2-4; see SI Fig.1 for detailed work flow). Segmented structures were assigned individual colours to differentiate them (as demonstrated in Fig.1b,d,f, and g). In Drishti, we use both 1D and 2D transfer functions with combinations of different illumination methods such as applying different opacity and light volumes per layer or segmented structures to revisualize the segmented volume data. Our method also allows each segmented layer of the original segmentation to be shown separately in the 2D transfer function window with its own voxel information (Fig.1g-h). The 2D transfer function then maps the voxel information of each layer to optical properties before applying different illumination methods. Voxel intensity per layer can be read separately or together using the transfer function editor.

**Figure 1.**
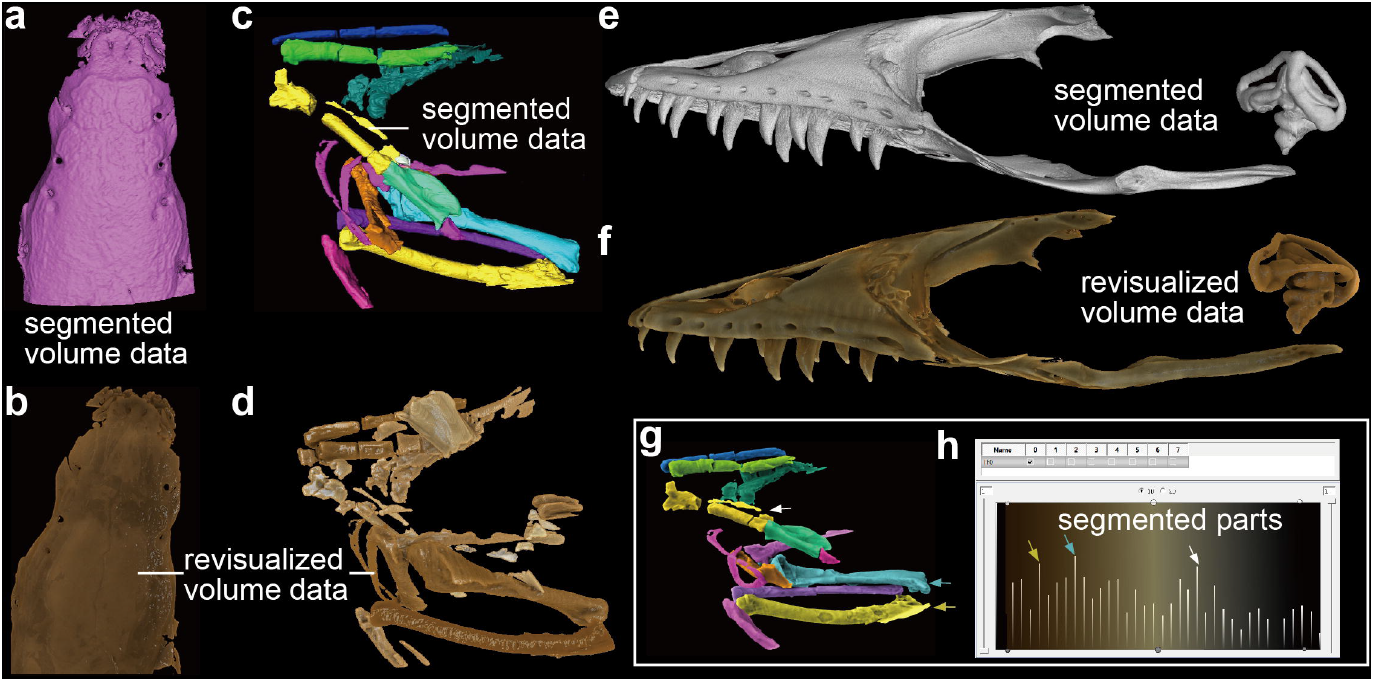
Segmented volume data from Mimics, Avizo and VG Studio and their corresponded revisualized segmented volume data from *Drishti*. **a** and **b,** Dorsal view of the braincase of living fish *Erofoichthys* IVPP OV2715 (**a,** Image obtained from Mimics for segmented volume; **b**, image obtained from *Drishti* after revisualising of segmented volume). **c** and **d**, Lateral view of segmented fossil bird *Linxiavis* IVPP V24116 (**c**, Image obtained from Avizo for segmented volume; **d**, image obtained from *Drishti* after revisualising of segmented volume). **e** and **f**, Lateral view of segmented lizard *Varanus indicus* (AMNH R58389) (**e,** Image obtained from VG Studio for segmented volume; **f**, image obtained from *Drishti* after revisualising of segmented volume). **g** and **h**, Demonstration of capability of assigning individual colours to differentiate segmented structures, allowing each segmented layer of the original segmentation to be shown separately in the 2D transfer function window with its own voxel information. Three pairs of coloured arrows indicate three separate revisualized segmented structures (**g)** and their corresponded counterparts in the 2D transfer function window (**h**).

Revisualized segmented volume data can be exported in three standard formats using *Drishti Import*: RAW, grayscale 8-bits unsigned image stacks and ITK MetaImage format, which are considered as standard formats that all image processing and 3D visualization software can read and process. Revisualized segmented volume can also be merged with original volume data for enhancement of a region-of-interest or segmented area which can validate the precision and accuracy of segmented results. To achieve this, two volumes, the revisualized segmented volume and the original tomogram need to be loaded together in *Drishti Render*, rendered then saved as an Extensible Markup Language data file (i.e. .xml). As demonstrated, the interchangeability between different image processing and 3D visualization software is achievable with this new revisualization method. In general, this method works for any software that allows DICOM as an output format. During the revisualization process, 2D and 3D images, movies and other visuals can also be generated. All revisualized volume data can be imported directly to *Drishti Prayog* and *Drishti VR* for instant interactive display.

Our revisualization method illuminates detailed information of segmented volume data through different combination of illumination algorithms and features. It utilizes mainstream image processing and 3D visualization software; maximizes the 3D visualized outcome of any segmented volume data while maintaining the uniformity and the interoperability between different software platforms and volumetric data formats. In addition, this method is the simplest way to break current restrictions and barriers due to compatibility, accessibility and affordability issues while maintaining the capability of producing more intuitive visual representations ensuring a more efficient collaboration between parties. By introducing this new revisualization method to the community will make examining current hypotheses or re-investigating published research findings are more achievable and feasible. This revisualization method is simple enough to be generally applied to any fields.

## Material and methods

### Segmented volume data for revisualization

Segmented volume data of the earliest tetrapodomorph, *Tungsenia paradoxa* (IVPP V10687). Original segmentation was done in Mimics^28^. Segmented braincase of *Erofoichthys* (IVPP OV2715) was scanned at IVPP. Both data were output as multiple zipped DICOM files from Mimics 18.0 (option: Export-Masks).

Segmented volume data of a Miocene bird *Linxiavis*^29^. Original segmentation was done in Avizo. This data was output as multiple DICOM directories from Avizo 9.0 (option: Export-DICOM).

Segmented volume data of lizard *Varanus indicus* (AMNH R58389) was done in VG Studio. This data was output as single DICOM directories from VG Studio 3.3 (option: Filter-Export- volume). The CT scanning of *Varanus indicus* was completed at the Microscopy and Imaging Facilities at the American Museum of Natural History (AMNH), with permissions from the Herptetology Department of the AMNH.

The above four experiments were then imported into *Drishti* Import for processing.

### Procession of segmented volume data

We have implemented a new ability to import multiple DICOM volume datasets and collapsing them into a single volume dataset. This was done in order to accommodate the segmentation results in the form of multiple DICOM volumes, each volume consisting of a single segmented structure. Each segmented structure is assigned a voxel value, thereby merging all the segmented datasets into a single volumetric dataset to be visualized.

We use two illumination methods in our revisualization experiments, the global parameters, which include the properties of the overall scene (e.g., lights, shading, camera position, projection type), and object parameters, which include the properties of the segmented object (e.g., colour and opacity). Global parameters were used to operate the scene of a 3D visualization. Different mixture of parameters were used in revisualized segmented volume data (see SI Figs. 2-4).

### Image analysis

Images of the revisualized volume data were generated in same orientation and scale in *Drishti* as in Mimics, Avizo and VG Studio. Image were analysed using ImageJ (https://imagej.nih.gov/ij/index.html) and ICY (http://icy.bioimageanalysis.org). Results were analysed use Microsoft Excel and displayed in the method and supplementary information. ImageJ (https://imagej.nih.gov/ij/index.html) and ICY (http://icy.bioimageanalysis.org/) were used to analysis images to demonstrate their properties (Fig. 2 and SI Figs. 2-4). Images were input into both ImageJ and ICY then transformed from RGB to 8-bits greyscale images for analysis. 2D histogram of pixel intensity, 3D surface plot and edges on those images were generated per image. The 2D histogram shows the distribution of grey values in that image. The x-axis represents the possible grey values and the y-axis shows the number of pixels found for each grey value. The 3D surface plot displays a three-dimensional graph of the intensities of pixels in a 8-bits greyscale colour image. Edges were calculated using Sobel Edge Detection algorithm ^30^. Results of these analysis are displayed here (Fig.2 and SI Figs.2-4).

**Figure 2.**
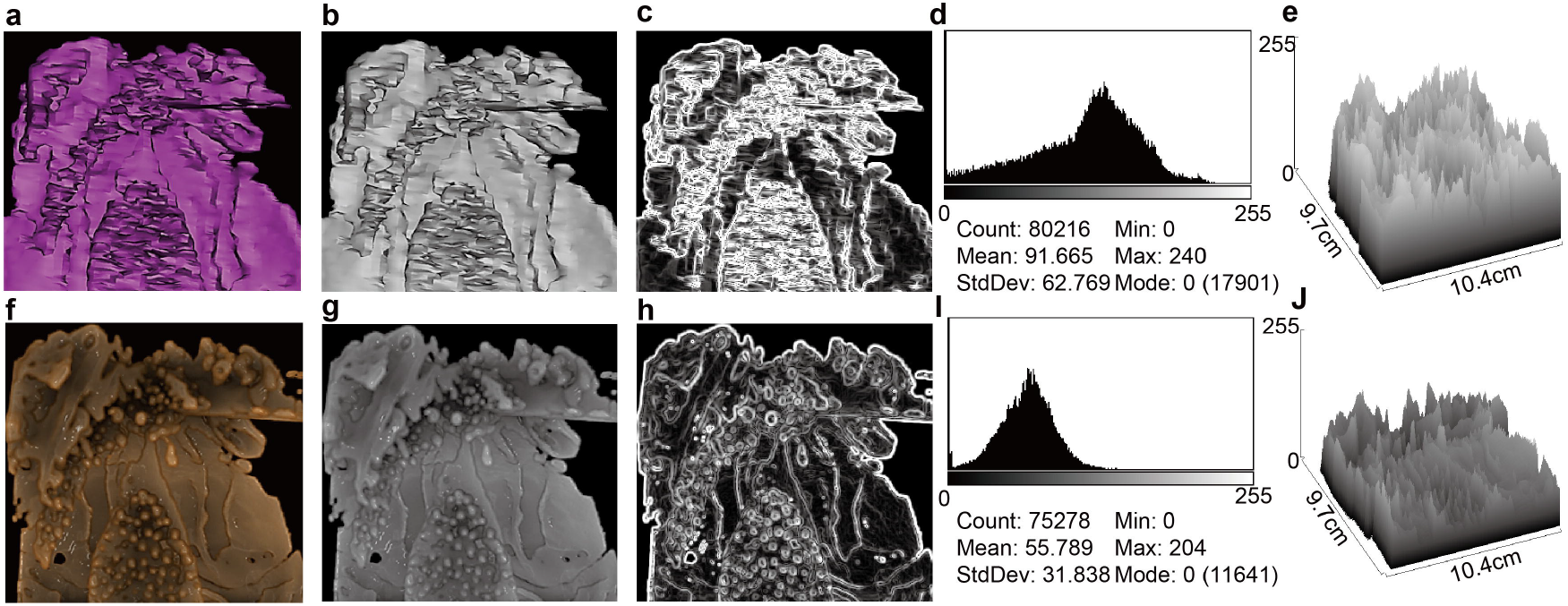
Comparison of segmented braincase of living fish *Erofoichthys* (IVPP OV2715) in Mimics and Dristhi. **a-c**, Images obtained from Mimics. **a**, Original image output from Mimics. **b**, 8-bits greyscale image of **a**. **c**, Edges in **b**. **d**, 2D histogram of **b**. **e**, 3D surface plot of **b**. **f-h**, Images obtained from *Drishti* after revisualization. **f**, Original image output from *Drishti* in the same orientation and scale of **a**. **g**, Revisualized result corresponds to **b**. **h**, Revisualized result corresponds to **c**. **i**, 2D histogram of **g. j**, 3D surface plot of **g**. Median Optical Density (OD) of **b** and **g** is 0.549 and 0.926 respectively, calculation carried out using the calibrated curve displayed in SI Fig. 8b.

The median optical density (OD) of each selected image (see SI Fig. 5) were measured in ImageJ for further analysis and displayed in SI Fig. 5. Calibration of optical density (SI Fig.8) was done using a Kodak No. 3 Calibrated Step Tablet scanned with an Epson Expression 1680 Professional scanner. The tablet has 21 steps with a density range of 0.05 to 3.05 OD (SI Fig. 8a). A weak correlation and positive linear relationship between images of the revisualized segmented volume using *Drishti* and images of the segmented volume in Mimics (SI Fig. 7).

## Data Availability

Step-table and calibrated curve for optical density and one movie for demonstration of the suggested workflow of revisualization are available through Figshare: https://figshare.com/s/350c258bc3d6b41313a1.

## Code Availability

Code is available at: https://github.com/nci/Drishti

## Acknowledgements

We thank Z.L. and H.Y. for sharing their segmented volume data, AMNH with permissions for using their data, T.S., M.Z. and G.C.Y. for their continuous support. This research was funded by the Strategic Priority Research Program of the Chinese Academy of Sciences (Grant No. XDB26000000) and the National Natural Science Foundation of China (41872023). Y.H. was supported by Postgraduate Research Scholarship at the Research School of Physics, Australian National University.

## Author Contributions

J.L. conceived the project. Y.H., A. J. and J.L. conducted functionality, unit and performance testing for *Drishti*. A.L. J.L. and Y.H. implemented the revisualization process and its corresponding illumination methods for DICOM directories. Y.H. and J.L. conducted image analysis and interpreted the results. All authors wrote the manuscript and contributed equally to this work.

## Competing Interests

The authors declare no competing interests.

## Supplementary information

Supplementary information is available to download here.

## Notes

### Competing Interest Statement

The authors have declared no competing interest.

