## Supplementary material for "A spark of 3D revisualization: new method for re-exploring segmented data": SI text

**This supplementary Information includes:**

Supplementary Notes

Supplementary Figures

### Supplementary Notes

#### *Drishti* v2.6.6

##### Installation

*Drishti* v2.6.6 is available to download from https://github.com/nci/*Drishti* under "Releases".

When you click, it will lead to a version-specific webpage where the.zip file is located at the bottom of the page, click *Drishti* v2.6.6.zip to download *Drishti*.

Source code and detailed information about this new release are also available on Github. Please note that *Drishti* v2.6.6 currently can only run on the Windows operating system. However, users can compile and install *Drishti* v2.6.6 for CentOS/Ubuntu. Sample compilation script for *nix systems is provided to users with the source code.

Once the download is finished, unzip the .zip file. Go to "bins" folder, then you should be able to run *Drishti* v2.6.6. *Drishti* is designed as a portable application thus making sure you know where the portable directory stores on your computer.

##### MIT License

Copyright (c) 2011-2020 National Computational Infrastructure, Australia.

### Supplementary Figures

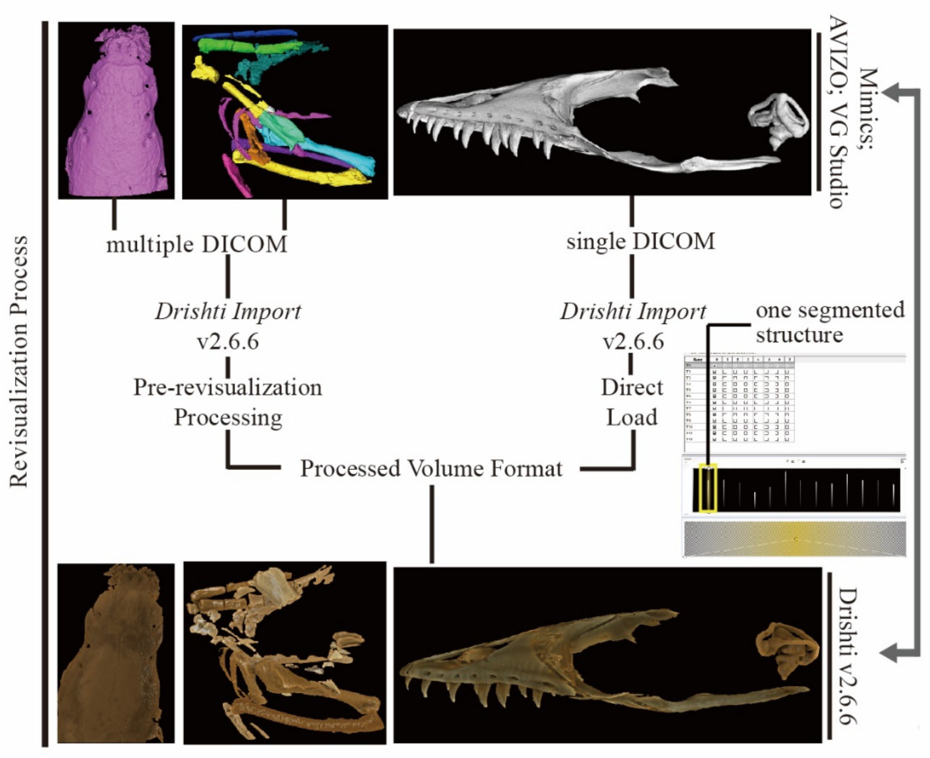


**Supplementary Figure 1. Workflow of revisualization process across three different popular image processing and 3D visualization software using the new version of the volume exploration and rendering software *Drishti* v2.6.6.**

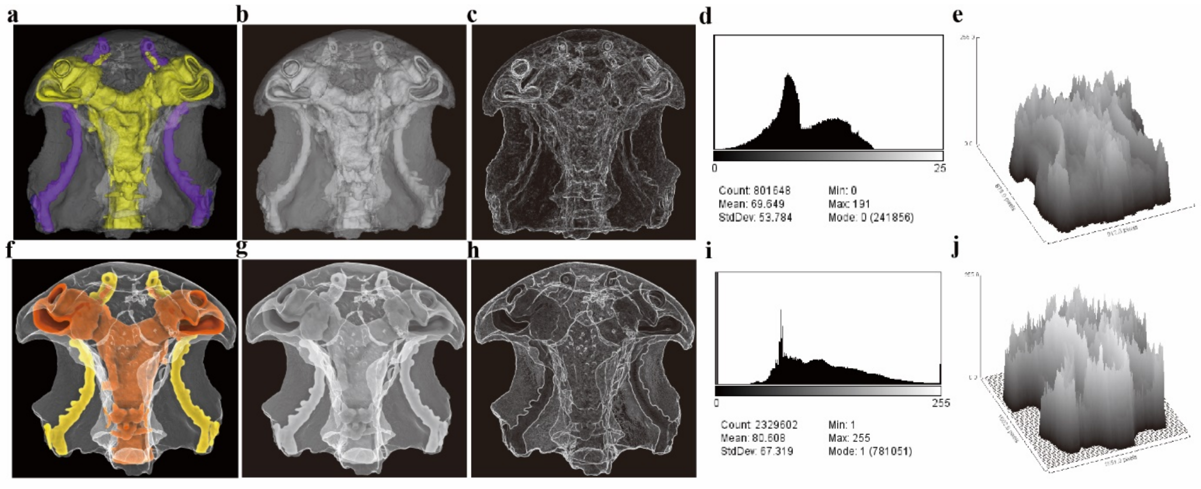


**Supplementary Figure 2. Ventral view of segmented braincase of fossil fish *Tungsenia* IVPP V10687. a-c**, Extracted images from Mimics. **a**, Original image output from Mimics. **b**, 8-bits greyscale image of **a**. **c**, Detected edges of **b**. **d**, 2D histogram of **b**. **e**, 3D surface plot of **b**. **f-h**, Extracted images from *Drishti* after revisualization. **f**, Original image output from *Drishti* in the same orientation and scale of **a**. **g**, Revisualized result corresponds to **b**. **h**, Revisualized result corresponds to **c**. **i**, 2D histogram of **g. j**, 3D surface plot of **g**.

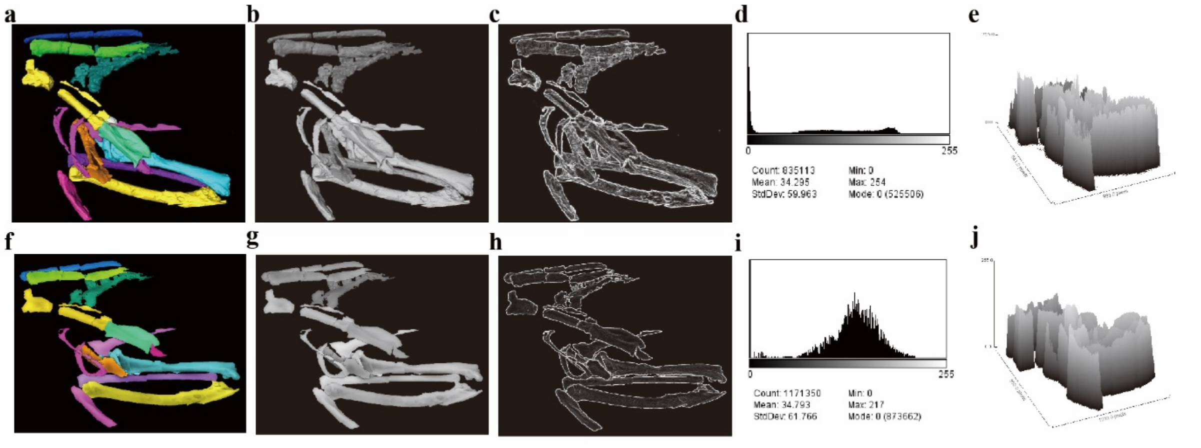


**Supplementary Figure 3. Lateral view of segmented fossil bird *Linxiavis* IVPP V24116. a-c**, Extracted images from Avizo. **a**, Original image output from Mimics. **b**, 8-bits greyscale image of **a**. **c**, Detected edges of **b**. **d**, 2D histogram of **b**. **e**, 3D surface plot of **b**. **f-h**, Extracted images from *Drishti* after revisualization. **f**, Original image output from *Drishti* in the same orientation and scale of **a**. **g**, Revisualized result corresponds to **b**. **h**, Revisualized result corresponds to **c**. **i**, 2D histogram of **g. j**, 3D surface plot of **g**.

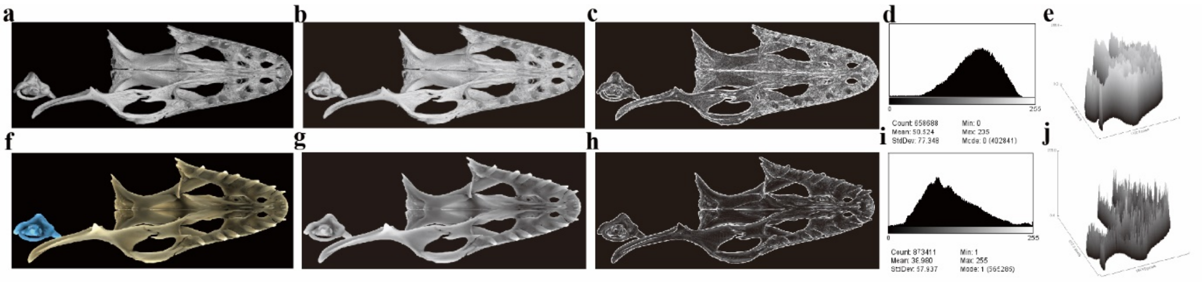


**Supplementary Figure 4. Ventral view of segmented upper jaw and inner ear of lizard Varanus indicus (AMNH R58389). a-c**, Extracted images from Mimics. **a**, Original image output from Mimics. **b**, 8-bits greyscale image of **a**. **c**, Detected edges of **b**. **d**, 2D histogram of **b**. **e**, 3D surface plot of **b**. **f-h**, Extracted images from *Drishti* after revisualization. **f**, Original image output from *Drishti* in the same orientation and scale of **a**. **g**, Revisualized result corresponds to **b**. **h**, Revisualized result corresponds to **c**. **i**, 2D histogram of **g. j**, 3D surface plot of **g**.

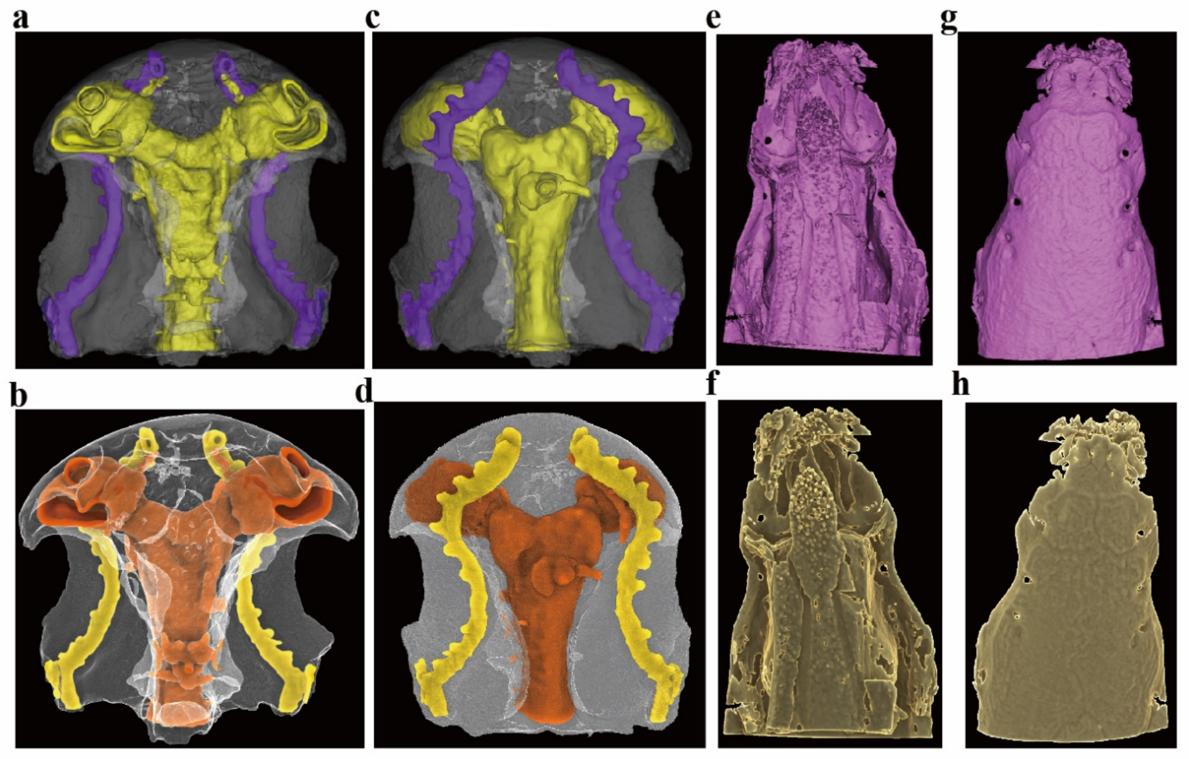


**Supplementary Figure 5. Images used for optical density measurements.** All images are from segmented volumes using Mimics and their corresponded revisualized segmented volume data using Drishti. **a-d**, Extracted images of *Tungsenia* IVPP V10687. **a-b**, Ventral view, from segmented volume (**a**) and revisualized segmented volume (**b**). **c-d**, Dorsal view, from segmented volume (**c**) and revisualized segmented volume (**d**). **e-h**, Extracted images of *Erofoichthys* IVPP OV2715. **e-f**, Ventral view, from segmented volume (**e**) and revisualized segmented volume (**f**). **g-h**, Dorsal view, from segmented volume (**g**) and revisualized segmented volume (**h**).

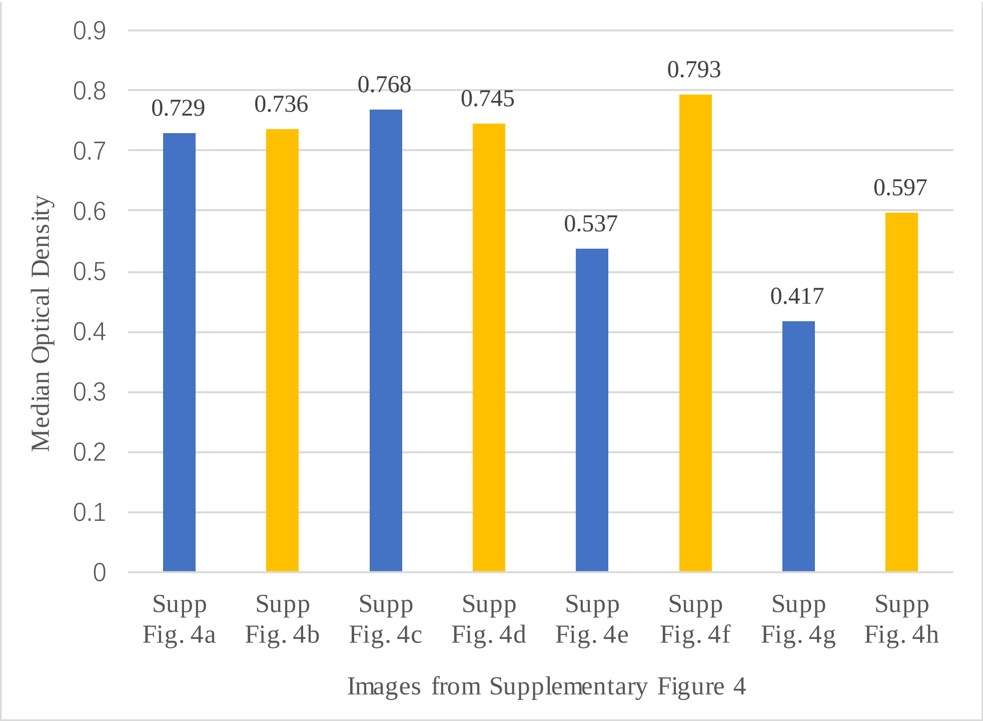


**Supplementary Figure 6.** Median optical density of images from Supplementary Fig.4. Calibration of optical density was completed using 8-bits 21 steps-table using ImageJ. Data analysis was carried out in Microsoft Excel.

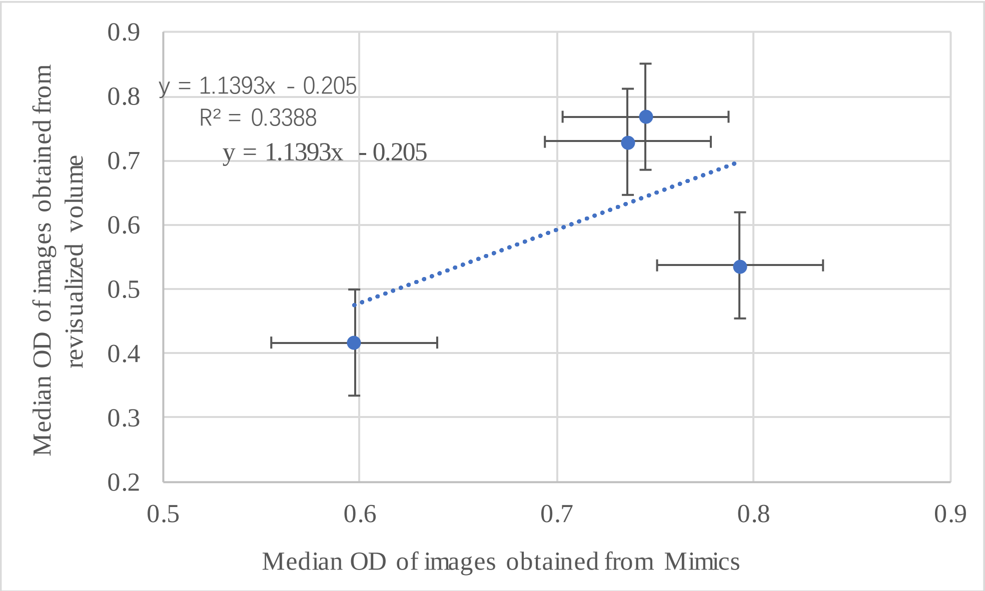


**Supplementary Figure 7.** Correlation between the median optical density of images in Supplementary Fig.4. X-axis represents the median optical density of images obtained from original segmented volume using Mimics (i.e. Supplementary Fig.4a,c,e,g). Y-axis represents the median optical density of images obtained from the revisualized volume using Drishti (i.e. Supplementary Fig. 4b,d,f,h). Analysis was carried out in Microsoft Excel.

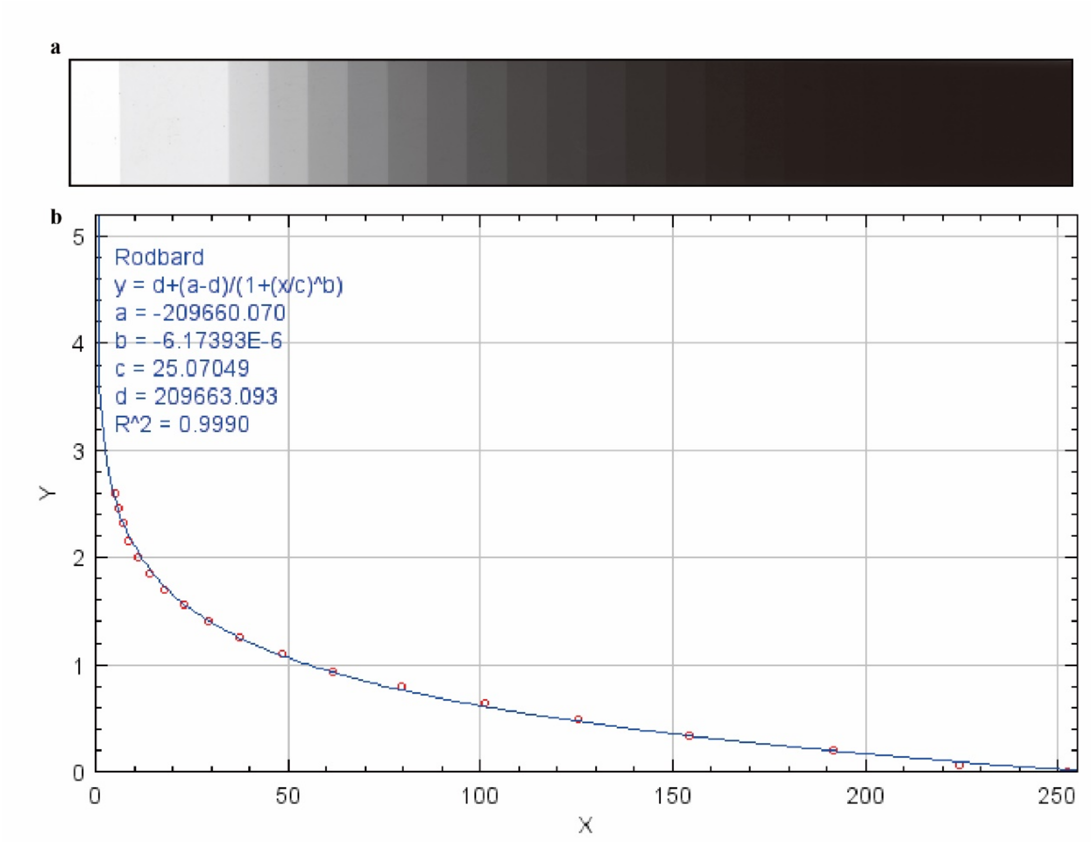


**Supplementary Figure 8. Step table and calibration curve used for this study.** a, step-table. b, calibrated Optical Density curve with the correlation coefficient R^2 equals to 0.9990.
